# Sensitivity Analysis of Biochemical Systems Using Bond Graphs

**DOI:** 10.1101/2023.04.04.535518

**Authors:** Peter J. Gawthrop, Michael Pan

## Abstract

The sensitivity of systems biology models to parameter variation can give insights into which parameters are most important for physiological function, and also direct efforts to estimate parameters. However, in general, kinetic models of biochemical systems do not remain thermodynamically consistent after perturbing parameters. To address this issue, we analyse the sensitivity of biological reaction networks in the context of a bond graph representation. We find that the parameter sensitivities can themselves be represented as bond graph components, mirroring potential mechanisms for controlling biochemistry. In particular a *sensitivity system* is derived which re-expresses parameter variation as additional system inputs. The sensitivity system is then linearised with respect to these new inputs to derive a linear system which can be used to give local sensitivity to parameters in terms of linear system properties such as gain and time constant. This linear system can also be used to find so-called sloppy parameters in biological models. We verify our approach using a model of the Pentose Phosphate Pathway, confirming the reactions and metabolites most essential to maintaining the function of the pathway.

## 1 Introduction

The *sensitivity* of a system is a quantitative measure of how much system outputs change in response to changes in parameters or inputs; a system that is insensitive is said to be *robust* with respect to those parameters, which are then refered to as *sloppy*. On the other hand, parameters associated with high sensitivity are often important for the function of a biological system, for example defining switch-like behaviours [1]. Sensitivity theory of dynamical systems and its application to engineering systems is well established and summarised in the textbooks [2, 3]. There are many applications of sensitivity methods to systems and control problems including: system optimisation, control system analysis [4, 5] controller tuning [6] and parameter estimation [7, 8].

The importance of sensitivity to the discipline of Systems Biology has long been recognised [9–12] and continues to be investigated [13–15]. In particular, Metabolic Control Analysis (MCA) [16–20] applies sensitivity methods to the analysis of chemical reaction networks in general and metabolism in particular. A thorough investigation into the links between MCA and more general sensitivity theory can be found in the works of Ingalls [21–24].

Because of the overlap between Engineering and Systems Biology, it is insightful to import methods from the former to the latter [25–27]. The bond graph technique provides one example of this methodological transfer, and has shown great potential in constraining models to be consistent with the laws of physics and thermodynamics [28, 29]. Bond graphs were introduced by Paynter [30, 31] to model the flow of *energy* though physical systems of interest to engineers and are described in several text books [32–35] and a tutorial for control engineers [36]. Bond graphs provide a systematic approach to elucidating the *analogies* between disparate physical domains based on energy and thus can also provide a foundation for Systems Biology. In particular, bond graphs were first used to model chemical reaction networks by Katchalsky and coworkers [37]. A detailed account is given by Oster et al. [38] and an overview of the approach is given by Perelson [39]. This work has been extended to Systems Biology [40–46]. A recent introduction and overview of the bond graph approach to systems biology is given by Gawthrop and Pan [28]. Sensitivity analysis is often based on analysing the ODE derived from a system rather than the system itself [12, 15]. In contrast, this paper analyses the system itself by extending the bond graph approach to sensitivity analysis, that has previously been developed for engineering systems [47–50], to biochemical systems. This approach has the advantage that the sensitivity is with respect to physically-meaningful parameters and both unperturbed and perturbed systems obey basic physical laws such as mass and energy conservation.

A key insight in sensitivity analysis is that the sensitivity properties of a system may be described by a related system: the *sensitivity system* [2, 48, 49, 51]. This notion has been extended in the context of bond graph models of engineering systems to the idea that a system represented by a bond graph has a sensitivity system also described by a bond graph [48, 49]. As will be shown, this property is inherited by biochemical systems.

Employing this sensitivity system in place of analysing the sensitivity of the underlying ODE has a number of advantages including: the variability is encapsulated by additional inputs rather than variable parameters; as the sensitivity system is itself a bond graph, the full range of bond graph tools can be applied including linearisation [41] and modular modelling [41, 44, 45].

There are two main categories of sensitivity analysis: local and global [12]. The former corresponds to small parameter variations and leads to linear dynamic systems; the latter corresponds to arbitrary parameter variations [12, 15] and is thus corresponds to nonlinear dynamic systems. In the context of this paper, the bond graph sensitivity system represents the nonlinear system and thus can be used for global analysis. As will be shown, this system can then be linearised and used for local sensitivity analysis. We focus on local sensitivity in this paper. Although the local approach is approximate, it is more computationally efficient than global methods and offers more insight into the effects of varying combinations of parameters. Indeed, the properties of the linearised system can be summarised by basic linear system properties such as gain and time constant; these can be derived without simulation. The variation of the local model properties with the point of linearisation can also give insight into system properties. The local approach can also be for initial investigations followed by targeted global analysis [52].

As mentioned above, parameter variation appears in the sensitivity system as additional inputs. Hence local sensitivity analysis, which linearises with respect to parameter variation, is closely related to linearisation with respect to inputs; this paper explicitly shows this relationship. Such linearisation has already been examined in the bond graph systems biology context [41, 53]. This paper looks at the approximation inherent in local analysis by comparing global with local results for a range of examples.

Sensitivity analysis can either be static and show how steady-state values of species depend on parameters, as in MCA, or be dynamic and show how the time trajectories of species depend on parameters. This paper looks at *dynamic* sensitivity analysis; this enables not only static values of sensitivity to be derived but also the sensitivity of the time course of species as parameters change. These sensitivity trajectories can be represented as a transfer function, the properties of which can be summarised in various ways according to the application. For example, the trajectory transfer function can be summarised by steady-state gain (also know as DC gain), initial response and time constant or, in the context of control theory, by a frequency response.

This paper extends bond graph based sensitivity analysis to biochemical systems. In particular, the bond graph of the sensitivity system of a biochemical system is derived; following [48], this involves ascertaining the sensitivity component associated with biochemical bond graph components. Biochemical bond graph components are different from those found in engineering systems and thus require the new methods derived in this paper. Biochemical systems form a subset of nonlinear systems with special properties; in particular, the mass-action equations result from the logarithmic nature of chemical potential [40, 54] and the exponential nature of the Marcelin – de Donder formulation of reaction flows [40, 55]. Moreover, the reaction components have two energy ports as compared to the single port of typical engineering resistive components. As will be shown, this leads to the reformulation of parametric variation in terms of modulating chemical potentials which act as additional system inputs.

Bond graph models of mass-action equations can be combined in a modular fashion to represent more complex reaction formulations such as enzyme catalysed reactions [28, 40, 41]; thus sensitivity of these more complex reaction formulations is within the scope of this paper.

It is well known that system dynamics can be insensitive to variation of parameters: this behaviour has been called *Sloppy Parameter Sensitivities* [56]. In particular, a quadratic cost function involving system trajectories has been formulated [56] to investigate this behaviour.

This paper draws explicit links with the Sloppy Parameter analysis of Gutenkunst et al. [56].

§ 2 gives the background in bond graph modelling required for the rest of the paper and introduces a set of normalised variables. § 3 introduces the bond graph sensitivity system appropriate to biochemical systems. § 4 shows how the nonlinear sensitivity system can be linearised. § 5 explores the relationship between the sensitivity system and the concept of sloppy parameter sensitivities. § 6 and § 7 look at two illustrative examples: a modulated enzyme-catalysed reaction and the Pentose Phosphate Pathway.

The Python code used to generate the Figures in this paper is available as Jupyter notebooks at https://github.com/gawthrop/Sensitivity23. Software tools for manipulating biochemical bond graphs are available in Python [57] at https://pypi.org/project/BondGraphTools/. and in Julia [58] at https://github.com/jedforrest/BondGraphs.jl. Both sets of tools are designed for modularity and scaleability and can generate both symbolic ODEs and numerical ODEs for simulation. The Python Control Systems Library, used for manipulating linearised systems, is available at https://pypi.org/project/control/.

## 2 Bond graph modelling of biochemical systems

This section summarises the material needed for the rest of the paper. § 2.1 summarises, in the form of a specific example, the basic ideas of modelling biochemical systems to be found in more detail in a recent paper [28]. § 2.2 introduces a normalisation approach to simplify the sensitivity analysis and § 2.3 considers the stoichiometry of bond graph models [28]. § 2.4 summarises the linearisation of biochemical systems [41].

### 2.1 Example: modelling the chemical reactions A⇄B⇄ C

This example considers the two chemical reactions 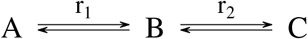 where the substance A is inter-converted with substance C via the intermediate substance B. Figure 1 gives the corresponding Bond Graph which consists of: the *components* **Ce** and **Re** which represent species and reactions respectively, the *bonds* ⇁ which carry energy in the form of chemical potential *μ* (J mol^−1^) and molar flow *v* (mol s^−1^) whose product *μ × v* has units of power (J s^−1^ or W) and the **0** junction which connects bonds in such a way that all impinging bonds carry the same potential. Thus, for example 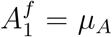 and *v*_*A*_ = *−v*_1_; similarly 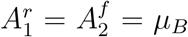 and *v*_*B*_ = *v*_1_− *v*_2_. Although not used in Figure 1, the dual **1** junction connects bonds in such a way that all impinging bonds carry the same flow. **Ce** components store, but do not dissipate, energy; **Re** components dissipate, but do not store, energy; bonds and junctions transmit energy without storage or dissipation.

**Figure 1.**
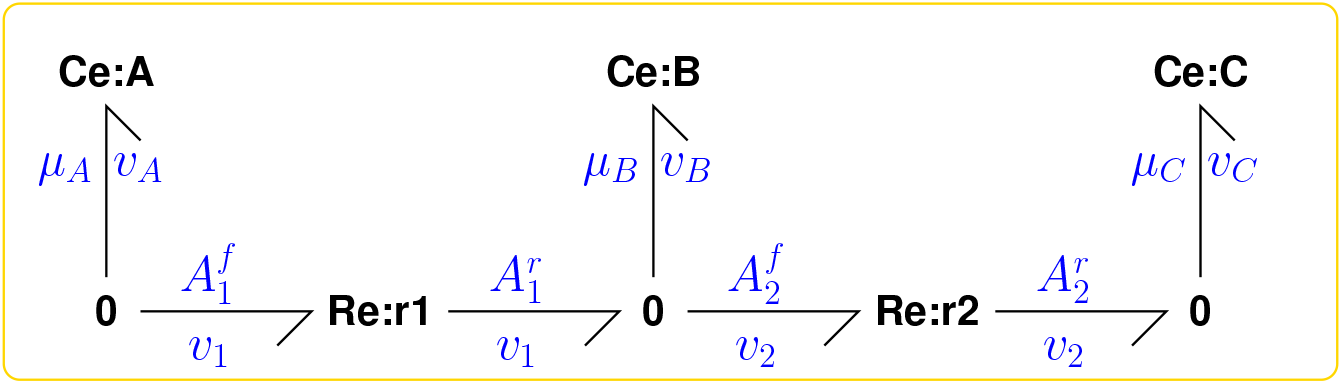
Example 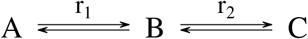. The three species A, B and C are represented by the three bond graph components **Ce**:**A, Ce**:**B** and **Ce**:**C** and the two reactions r_1_ and r_2_ by the two bond graph components **Re**:**r1** and **Re**:**r2**. The bonds ⇁ carry the chemical potential μ, or affinities comprising sums of potentials, and molar flow *v*, and thus energy flows, between components. The **0** junctions combine energy flows in such a way the all impinging bonds carry the same potential.

The **Ce**:**A** component has the constitutive equations

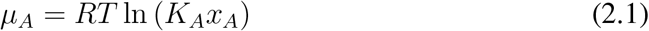

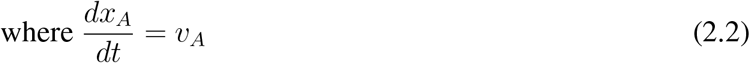

Where the time derivative in (2.2) means that this component is dynamic. R = 8.314 J mol^−1^ K^−1^ is the Universal Gas Constant and T (K) the absolute temperature. *μ*_*A*_ (J mol^−1^) is the chemical potential of substance A, *x*_*A*_ (mol) the amount of substance A and *v*_*A*_ (mol s^−1^) the molar flow of substance A. As discussed by Gawthrop and Pan [28] § 3.2, *K*_*A*_ (mol^−1^) is given by

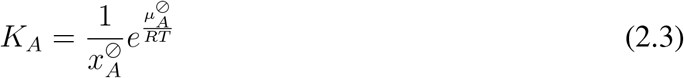

where 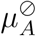 is the chemical potential of substance A when 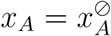.

In contrast, the **Re**:**r1** component is static with constitutive equation

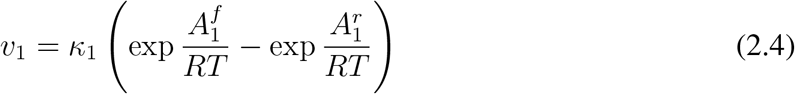

Where 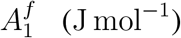 and 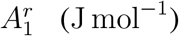are the forward and reverse affinities associated with reaction r_1_ and *v*_1_ (mol s^−1^) the molar flow associated with reaction r_1_. Analogous equations apply to the **Ce** components B and C, and the **Re** component r_2_.

In this simple example, these equations yield the well-known law of mass action

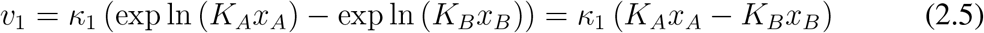

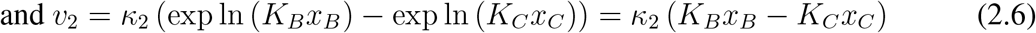

More complex biochemical system are needed to reveal the full power of the bond graph approach. For example detailed balance and Wegscheider conditions are automatically satisfied [28, 59].

As discussed previously [28, 41], an open system can be obtained from a closed system by replacing dynamic **Ce** components by *chemostats* [60] where the ODE (2.2) is replaced by

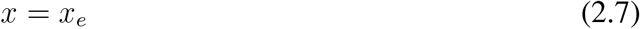

This example is continued throughout the paper as an open system with both **Ce**:**A** and **Ce**:**C** chemostats. This corresponds to the situation where substrate A and product C are kept at constant amounts.

### 2.2 Normalisation

It is convenient to re-express chemical potential *μ* (J mol^−1^) in non-dimensional form *ϕ* ^1^where

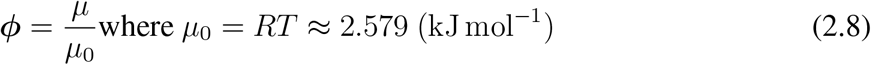

For this section, we derive numerical values assuming that T = 37°C ≈ 310 K. To re-express molar flow *v* (mol s^−1^) in non-dimensional form ***f***, define a normalising power *P*_0_ (W). Choosing *P*_0_ = 1 mW gives

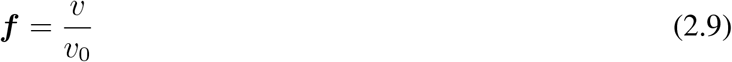

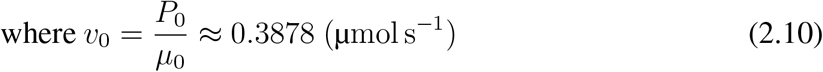

The amount *x* can be re-expressed as the normalised variable ***x***

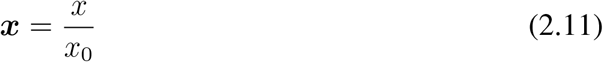

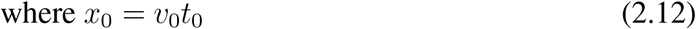

where *t*_0_ (s) is a normalising time unit and *v*_0_ is given by (2.10). We can also define the normalised time as ***t*** = *t*/*t*_0_. Thus, the derivative 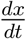 of the amount *x* can be re-expressed as the normalised variable 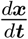 :

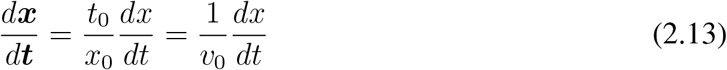

Using normalised variables, the **Ce** constitutive relation (2.1) becomes:

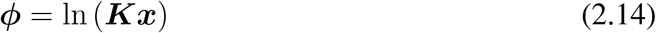

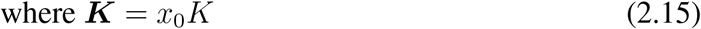

and the **Re** constitutive relation (2.4) becomes:

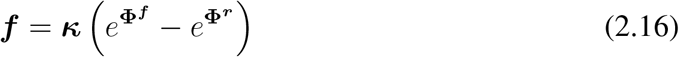

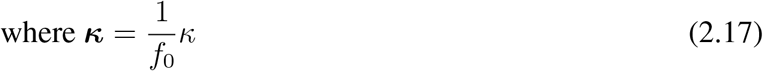

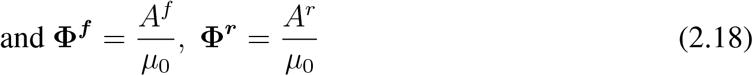

An alternative set of units uses the Faraday constant *F ≈* 96.49 (kC mol^−1^) to replace chemical potential *μ* (J mol^−1^) by 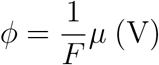 and molar flow *v* (mol s^−1^) by *f* = *Fv* (A) [28, 61]. In this case the normalised variables can be re-expressed as:

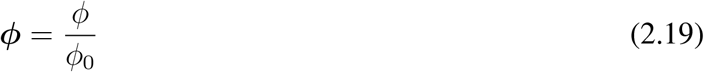

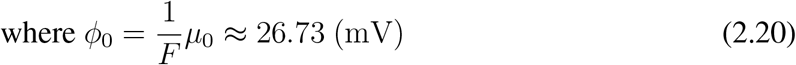

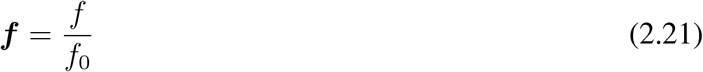

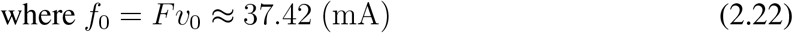

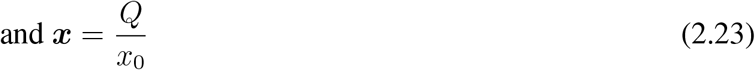

where *x*_0_ (C) is the amount expressed in Coulombs.

### 2.3 Stoichiometry

As discussed previously [28, 40, 45] the stoichiometric representation of a biochemical system can be derived directly from the bond graph. In normalised form, the stoichiometric representation is:

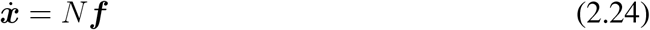

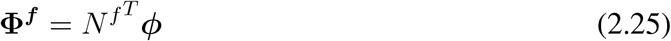

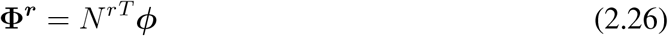

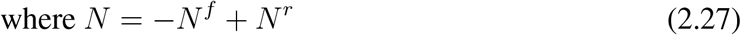

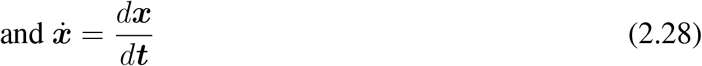

In the context of the example of § 2.1:

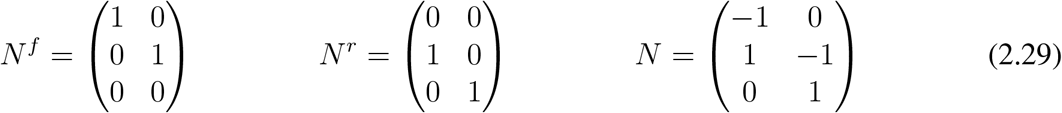

where

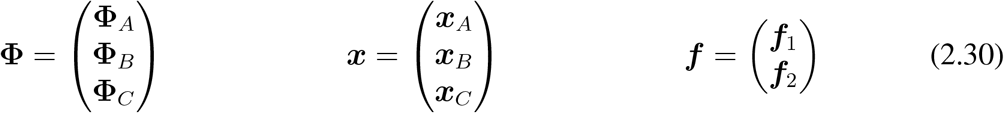

As discussed on § 2.1, an open system can be obtained from a closed system by replacing dynamic **Ce** components by *chemostats*. In stoichiometric terms, Equation (2.24) is replaced by

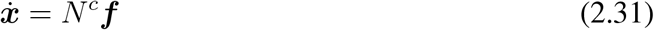

where the chemodynamic stoichiometric matrix *N*^*c*^ is identical to the stoichiometric matrix N except that rows corresponding to chemostats are set to zero [28, 41].

In the context of the example of § 2.1, choosing both **Ce**:**A** and **Ce**:**C** to be chemostats gives:

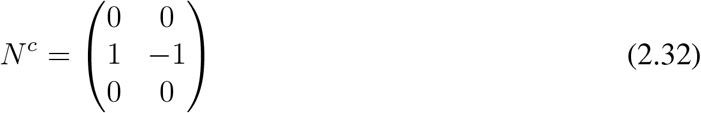

### 2.4 Linearisation

Linearisation in the context of bond graph models of engineering systems is discussed by Karnopp [62], who showed that the linearised system could also be represented as a bond graph. Linearisation in the context of bond graph models of biochemical systems is presented by Gawthrop and Crampin [41]. A key feature of biochemical systems is the presence of *conserved moieties* which lead to constraints on system states [40, § 3(b)] which in turn leads to reduced-order dynamical equations [40, § 3(c)]. If such state constraints are not explicitly accounted for, linearisation leads to systems with uncontrollable or unobservable states [63, 64] which appear as cancelling pole/zero pairs in the corresponding transfer functions. This paper uses the approach of Gawthrop and Crampin [41, § 4.2] to avoid this issue.

In particular, the linearised system is given in standard state-space form as:

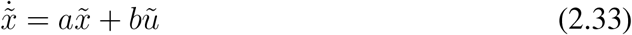

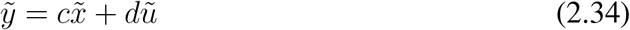

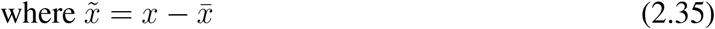

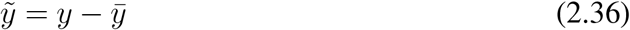

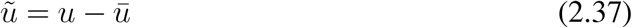

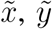 and 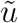 are the deviation of the state *x*, the output *y* and the input *u* from the steady state values 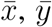 and 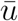 and *a, b, c* and *d* are matrices. The number of states, inputs and outputs are denoted *n*_*x*_, *n*_*u*_ and *n*_*y*_ respectively.

This linear system has the corresponding *n*_*y*_ *× n*_*u*_ transfer function matrix *G*(*s*) relating the *n*_*u*_ inputs to the *n*_*y*_ outputs given in terms of the Laplace variable s as:

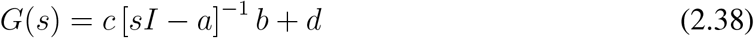

where *I* is the *n*_*x*_ *× n*_*x*_ unit matrix.

In engineering system analysis [5], it is standard practice to characterise the time (*t*) response of a linear system in terms of the system *unit step response*. Because there are *n*_*y*_ outputs and *n*_*u*_ inputs, the unit step response *g*_*ij*_(*t*) of the *i*th output with respect to the *j*th input is defined as the time response 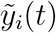of the *i* output when the initial state 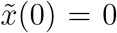 and each input 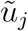is the unit step function *U*(t) where

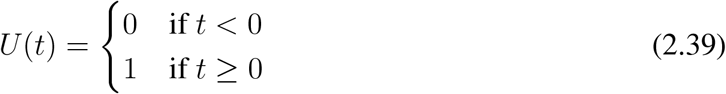

These individual step responses can be combined into the *n*_*y*_ *× n*_*u*_ matrix *g*(*t*) where the *ij*th element is *g*_*ij*_(*t*). In particular, if the vector of *n*_*u*_ inputs 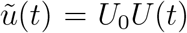 where the *j*th element of the vector *U*_0_ is the amplitude of the *j*th step input, then

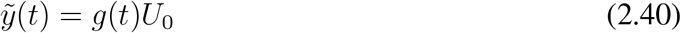

Moreover, the transfer function *G*(*s*) is the Laplace transform of the impulse response *g*′(*t*) = *dg*/*dt* and thus the Laplace transform of the the step response *g*(*t*) is *G*(*s*)/*s*. It follows from the final and initial value theorems of the Laplace transform [5] that the time and Laplace domains are linked by:

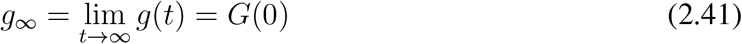

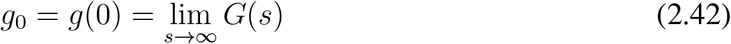

Equation (2.41) gives a convenient method for computing steady-state values from the transfer function and Equation (2.42) gives a convenient method for computing the initial response of the linearised system.

Although the actual sensitivity system response may be complicated, it is useful to provide an indicator *τ* of the overall time scale. In the case where *g*_0_ ≠ 0, the response is instantaneous and hence *τ* = 0. When *g*_0_ = 0, a first-order transfer function is of the form:

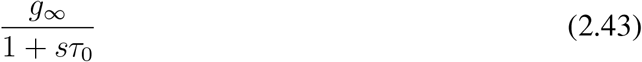

where *τ*_0_ is the system time constant. In this case it is convenient to use *τ* = *τ*_0_. In the general case, there are many possible time constants to choose. Here, we use *balanced order reduction* [65] to reduce the system order to unity and select the corresponding time constant as in the first order case.

The three numbers *g*_0_, *g*_*∞*_ and *τ* provide a convenient summary of the dynamic properties of the linearised system (2.33) – (2.34). In the control literature, *g*_*∞*_ is called the *DC gain* or *steady-state* gain of the linearised system (2.33) – (2.34).

## 3 The Sensitivity System

This section shows how the bond graph corresponding to the sensitivity system can be derived from the bond graph of the biochemical system itself. There are two approaches considered. § 3.1 shows how system components with variable parameters can be replaced by a sensitivity component represented itself by a bond graph thus giving an explicit bond graph for the sensitivity system. An alternative explored in § 3.2 is to derive the stoichiometric matrix of the bond graph biochemical system model, augment this matrix to give the stoichiometric matrix of the sensitivity system and then construct the corresponding bond graph. These conversions from stoichiometric matrix to bond graph and from bond graph to stoichiometric matrix are based on earlier work [45].

### 3.1 The sensitivity bond graph

The key idea in this paper is to replace each *parametric perturbation* by an *equivalent chemostat* (§ 2.1) and thus each bond graph component by a corresponding sensitivity bond graph component. This leads to:

1. a (nonlinear) sensitivity system represented by a bond graph for any biochemical system represented by a bond graph;
2. a (linear) local sensitivity system via linearisation [41] of the sensitivity system and
3. a stoichiometric interpretation of the sensitivity system.

#### 3.1.1 The Ce sensitivity component

The constitutive relation (2.14) of the normalised **Ce** component has a single parameter ***K***. For a given substance, ***K*** is dependent on temperature.

The perturbation in this parameter is modelled using the multiplicative variable λ so that the constitutive relation (2.14) is replaced by:

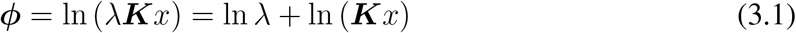

Thus the effect of λ can be thought of as adding a second chemostat **Ce** component in such a way that the normalised potential 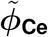 adds to the normalised potential *ϕ*_0_ of the original component to give

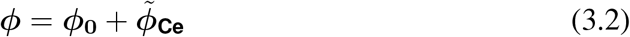

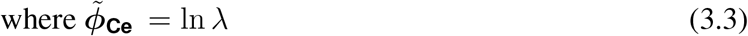

Figure 2(a) gives the bond graph interpretation.

**Figure 2.**
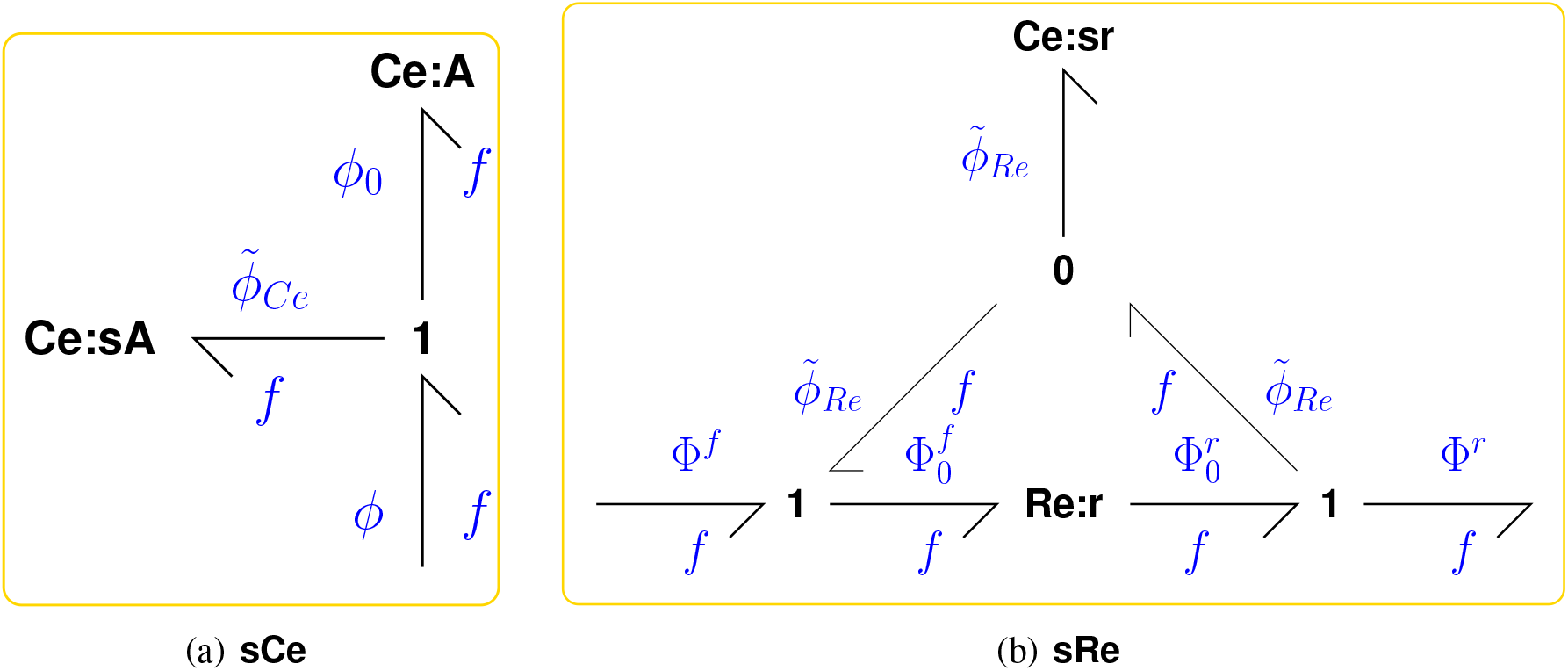
Sensitivity components (a) **sCe**, the sensitivity **Ce** component, comprises the original **Ce** component **Ce**:**A** with normalised potential *ϕ*_0_ and an additional chemostat **Ce**:**sA** with normalised potential 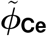 connected by a **1** junction so that the overall normalised potential is 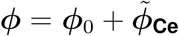. (b) **sRe**, the sensitivity **Re** component, comprises the original **Re** component **Re**:**r** with normalised forward and reverse potentials 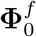 and 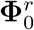 and an additional chemostat **Ce**:**sr** with normalised potential 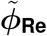 connected by **1** junctions so that the overall normalised forward and reverse potentials are 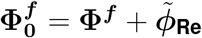 and 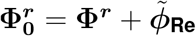.

##### The chemostat sensitivity component

In bond graph terms, *chemostats* are **Ce** components with a fixed state *x* = *x*_0_. As discussed in § 2.1 they represent substances such as metabolites which are external to the system. In biochemical terms, *x*_0_ can vary if the amount of the external substance changes. In this case, the fixed state *x*_0_ can also be perturbed using the multiplicative parameter *λ*:

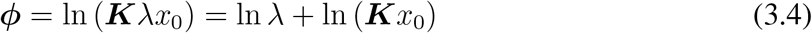

Comparing Equation (3.4) with equation (3.1) it follows that a perturbation in the chemostat state *x*_0_ can be modelled in the same way as a perturbation in ***K***.

#### 3.1.3 The Re sensitivity component

The constitutive relation (2.16) of the normalised **Re** component has a single parameter *κ*. In biochemical terms, *κ* can vary due to the amount of enzyme and the effect of enzyme inhibitors. The perturbation in the parameter *κ* is modelled using the multiplicative variable λ so that the constitutive relation (2.16) is replaced by:

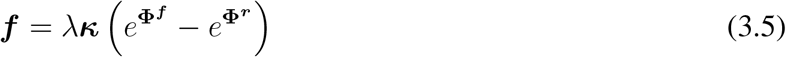

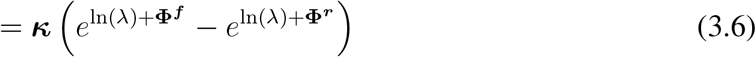

Thus the effect of *λ* can be thought of as adding a chemostat **Ce** component in such a way that the normalised potential 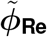 adds to the normalised forward and reverse reaction potentials **Φ**^***f***^ and **Φ**^***r***^ to give the normalised forward and reverse reaction potentials 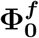 and 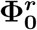 at the original

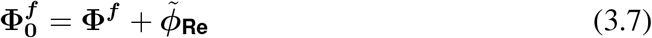

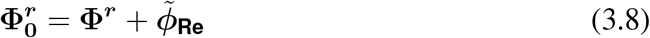

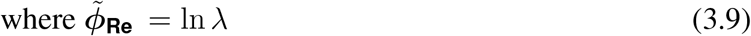

Figure 2(b) gives the bond graph interpretation.

Thus, for both the **Ce** and **Re** components, the perturbation λ is modelled as an additional chemostat with normalised potential 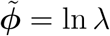.

#### 3.1.4 Example A ⇄B ⇄C

The sensitivity system of Figure 3 gives the normalised flows ***f***_1_ and ***f***_2_ through the reaction components **Re**:**r**_**1**_ and **Re**:**r**_**2**_ as:

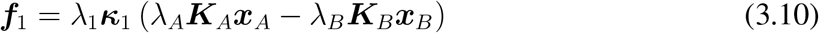

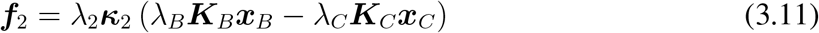

**Figure 3.**
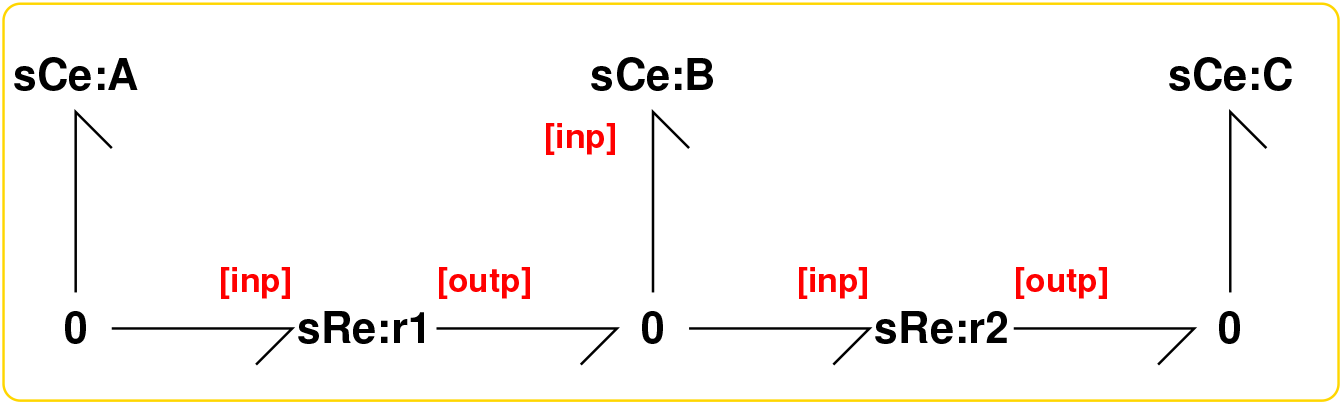
Sensitivity Example A ⇄B ⇄C. This figure is similar to Figure 1 except that the **Ce** and **Re** components have been replaced by the **sCe** and **sRe** components of Figure 2. Thus the additional **Ce** and **Re** components of Figure 2 are implicitly included within each sensitivity component. As discussed in § 2.1, the system is open and **Ce**:**A** and **Ce**:**C** are chemostats; the corresponding sensitivity components are modelled in § 3.1.2.

Note that if each *λ* = 1, the two flows correspond to the nominal system. The steady state flows correspond to 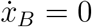; that is ***f***_1_ = ***f***_2_ and thus:

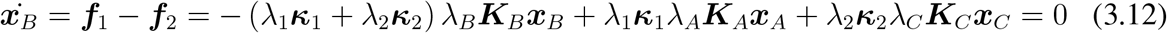

The steady-state value 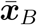 of ***x***_*B*_ is then given by:

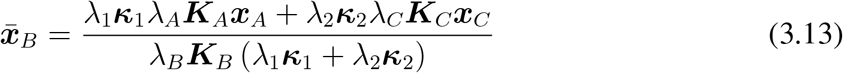

Again, the nominal steady state is obtained by setting each *λ* = 1.

### 3.2 A stoichiometric interpretation

The bond graph interpretation of the sensitivity system given in § 3.1 is conceptually useful and also provides a method for computer generation of the sensitivity system equations. This section presents an alternative approach to generating the sensitivity system equations from the bond graph using the stoichiometric representation outlined in § 2.3. For simplicity, the equations are generated for the sensitivity of every component.

The stoichiometry of the bond graph was considered in § 2.3. As discussed in § 3.1, the sensitivity bond graph consists of the original bond graph with extra **Ce** components acting as chemostats connected to the original **Ce** and **Re** components as indicated in Figure 2. These connections, together with those of the original bond graph, define the stoichiometry of the sensitivity bond graph. Following § 2.3, the equations for the closed sensitivity system where all chemostats are regarded as ordinary **Ce** components is considered first. Assuming that a sensitivity component is appended for every **Ce** component and every **Re** component, the sensitivity system state ***x***_*s*_ can be written as:

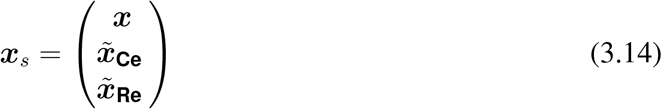

where *x* is the original system state and 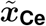 and 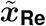 are the the states of the additional sensitivity **Ce** components associated with each **Ce** and **Re** components respectively.

Noting from Figure 2(a) that the sensitivity components associated with the **Ce** components share the same flows as the **Ce** components themselves, it follows from Equation (2.31) that, in the case of a closed system:

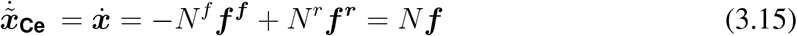

Further, noting from Figure 2(b) that the sensitivity components associated with the **Ce** components share the same forward (***f***^***f***^) and reverse (***f***^***r***^) flows as the **Re** components themselves, it follows that:

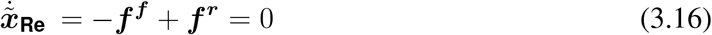

From § 2.3, it follows that the stoichiometry of the sensitivity system corresponding to the open system with 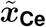 and 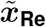 as chemostats is given in terms of the stoichiometry (Equations 2.31 – 2.26) by:

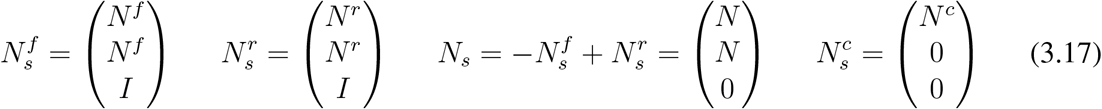

and

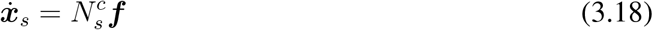

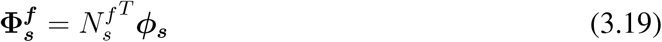

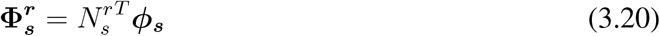

## 4 Linearisation of the Sensitivity System

§ 3 showed how the bond graph of the sensitivity system can be derived from the bond graph of the system itself; in general, this sensitivity system is nonlinear. A feature of the sensitivity system is that each *parameter variation* is replaced by a *chemostat* representing a system *input*. As discussed in § 2.4, nonlinear biochemical systems can be linearised. In the case of the non-linear sensitivity system of § 3 the inputs are the parameter variables *λ* (represented by chemostats) and system flows were chosen to be the outputs. Hence the sensitivity system can be linearised about a steady-state to give a linear system of the form:

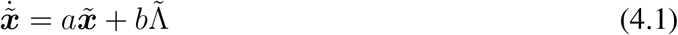

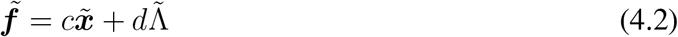

where the column vector 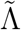 contains all of the sensitivity deviation variables 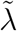. Comparison with Equations (2.33) and (2.34) shows that 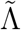 has the same role as the system input, and 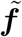 the same role as the system output, in the linearised system.

As discussed in § 2.4, the three numbers *g*_0_, *g*_*∞*_ and *τ* provide a convenient summary of the dynamic properties of the linearised sensitivity system (4.1) – (4.2). In particular, *g*_*∞*_ is the *DC gain* or *steady-state* gain of the linearised sensitivity system (4.1) – (4.2) and g_0_ and τ represent the initial response and time-scale respectively.

Using the result of Karnopp [62] noted in § 2.4, it follows that the linearised sensitivity system also has a bond graph representation but with linear components.

### 4.1 Example A ⇄B⇄ C

As an illustrative example, this section continues the example of § 2.1 for the particular set of parameters:

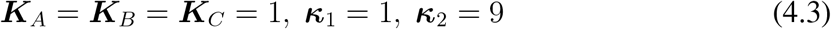

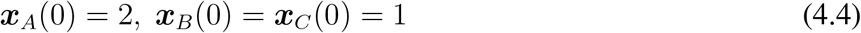

Using the explicit calculations of Appendix A, the dynamic linearisation of § 2.4 gives the transfer function:

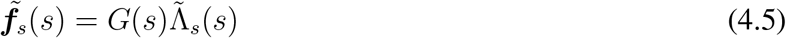

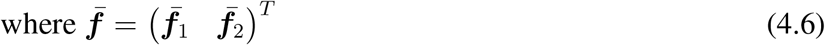

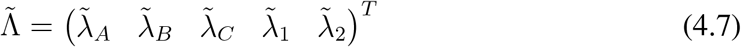

where 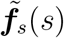 and 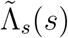 are the Laplace transforms of the flows 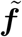 and the incremental sensitivity parameters 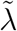 and *G*(*s*) is the transfer function. In this case:

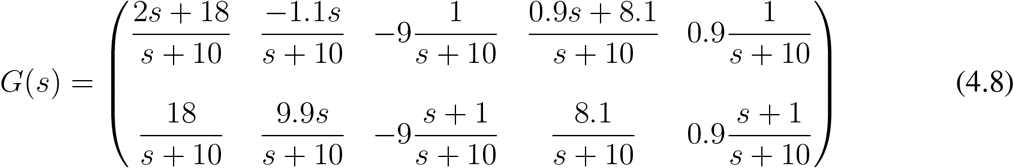

Using Equation (2.41) of § 2.4 the steady-state gains relating the two flows to the five parameters are:

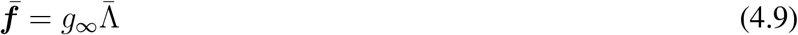

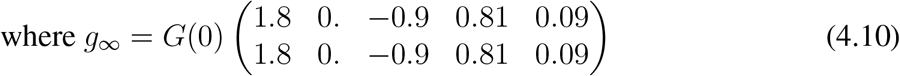

Using Equation (2.42) of § 2.4, the initial response is:

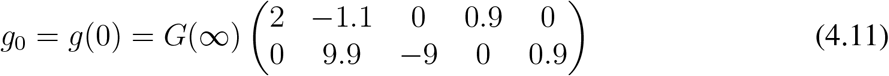

The five figures 4(a)–4(e) correspond to the five columns of *g*(*t*). To emphasise that the linearised response is a *linear* function of the perturbation amplitude, the figures show the perturbation step response 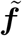 normalised by the perturbation step amplitude 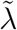. In each case, the normalised response of the flow through **Re** r1 and **Re**:**r2** to a 10% step change in the corresponding parameter is shown as dashed lines and compared to the unit step response of the corresponding transfer function element of *G*(*s*) together with the steady-stage gain *g*_*∞*_. The initial and final values correspond to the elements of the matrices (4.11) and (4.10) respectively as indexed by (4.6) and (4.7).

**Figure 4.**
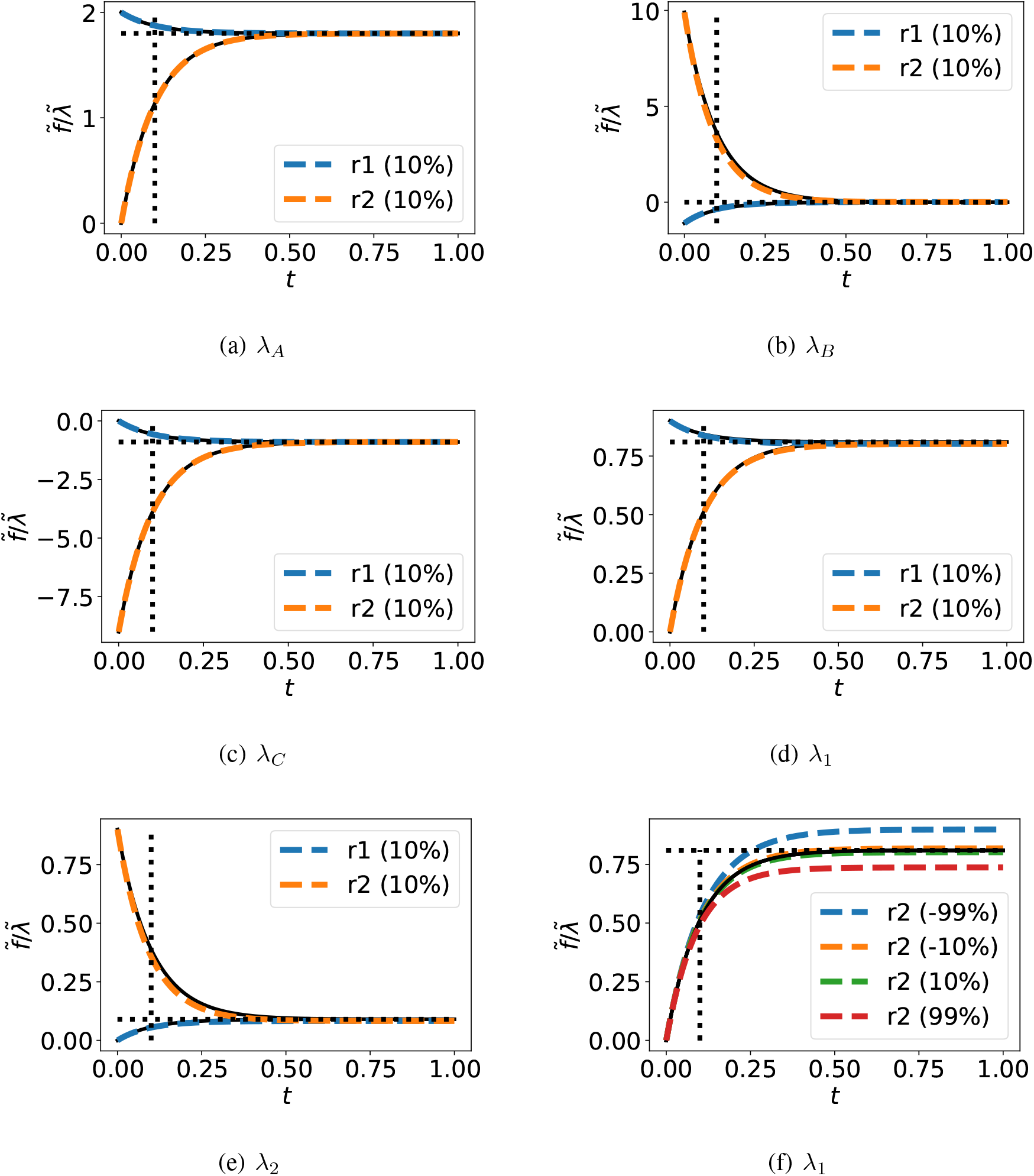
Illustrative example: the linearisation approximation. The normalised (with respect to perturbation 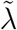 – see text) temporal responses of the flows through **Re** r1 and **Re**:**r2** to perturbations in (a) ***K***_*A*_; (b) ***K***_*B*_; (c) ***K***_*C*_; (d) ***κ***_1_; (e) ***κ***_2_ represented by the corresponding perturbation parameters *λ*_*A*_, *λ*_*B*_, *λ*_*C*_, *λ*_1_ and *λ*_2_ respectively as in Equations (4.5) – (4.7). Parameters are perturbed using a 10% step change in the perturbation parameter *λ*. The black line is the normalised response of the linearised sensitivity system to the step change in the perturbation parameter *λ* (i.e. the unit step response), the horizontal dashed black line the corresponding DC gain g_*∞*_ and the vertical dashed black line the corresponding time-constant *τ*. The coloured dashed lines show the response of the simulated (nonlinear) sensitivity system to the same perturbation in *λ*. (f) As (d) but the flow through **Re**:**r2** is shown for a *±*10% and *±*99% step change in the perturbation parameter *λ*_1_. In each case, the response of the linearised sensitivity system is an approximation to the response of the (nonlinear) sensitivity system itself.

The nonlinear response is close to that of the linearised response in each case. Figure 4(f) indicates how the discrepancy changes with parameter variations of *±*10% and *±*99%.

## 5 Sloppy Parameter Sensitivities

In their seminal paper, Gutenkunst et al. [56] introduced the concept of *Sloppy Parameter Sensitivities* in the context of fitting parameters to experimental data. Following standard system identification practice [8], their method is based on a quadratic cost function of the variation of the the system time response as parameters vary. This gives rise to a data-dependent Hessian matrix; the eigenvalues and eigenvectors of this matrix determine parameter sensitivity.

This section looks at how this concept applies in the context of the linear time responses of the linearised sensitivity model of § 4. As discussed in § 2.4, the linearised sensitivity system of § 4 can be summarised in terms of the two parameters: steady-state gain g_*∞*_ and time constant *τ* ; these two parameters can be deduced from the model itself *without* requiring simulation. In contrast, the sloppy parameter approach [56] is data based. Therefore the relationship between the two approaches to characterising model sensitivity to parameter variation is via the simulation of a model to generate data, even though such simulation is not required to generate *g*_*∞*_ and *τ*. As an illustrative example, the sensitivity of internal flows through reactions and external flows associated with chemostats are considered here. In particular, define the quadratic function *Q* of normalised system flow variation 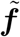:

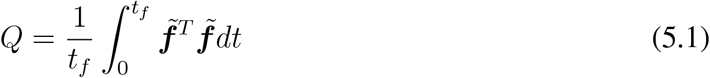

Using Equation (4.5), it follows that the cost function *Q* may be rewritten explicitly in terms of the parameter variation vector 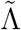 as

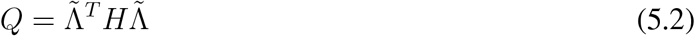

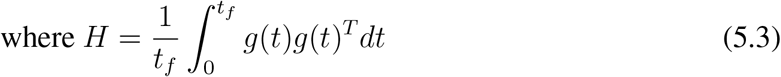

The *n*_Λ_ *× n*_Λ_ matrix *H* is positive semi-definite and therefore has the eigenvalue decomposition:

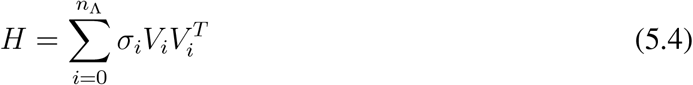

where *σ*_*i*_ *≥* 0 is the ith eigenvalue and the *n*_Λ_ column eigenvectors *V*_*i*_ are orthogonal and have unit length. Without loss of generality, assume that the eigenvectors are sorted with the largest first and the smallest last. The basic idea of sloppy parameter sensitivities [56] is that *small* eigenvalues *σ*_*i*_ correspond to directions (indicated by *V*_*i*_) in parameter space where the effect of parameter variations is small – these parameter combinations are termed *sloppy*. Conversely, *large* eigenvalues *σ*_*i*_ correspond to directions where the effect of parameter variations is large – these parameter combinations are termed *stiff*.

In the special case that the final time *t*_*f*_ is large, *H* can be approximated in terms of the steady state time response *g*_*∞*_ :

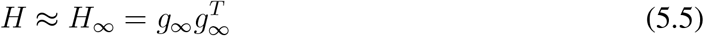

In the special case of a single output, *g*_*∞*_ is a vector and *H*_*∞*_ has rank 1 and the matrix *H*_*∞*_ has the decomposition

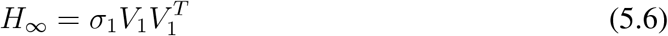

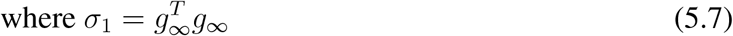

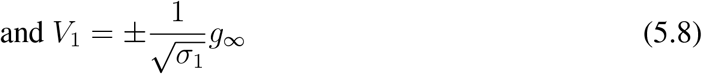

thus *n*_Λ_ − 1 eigenvalues are zero. The corresponding cost function is defined as:

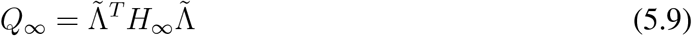

### 5.1 The two-parameter case

Because *Q* is a quadratic function of the sensitivity parameter vector Λ each constant value of *Q* corresponds to an ellipsoid in *n*_*λ*_-dimensional space. In particular, in two-dimensional parameter space, this eigen-decomposition gives elliptical contours of *Q* = 1 with minor axis corresponding to 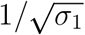 and major axis corresponding to 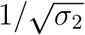 where *σ*_1_ ≥ *σ*_2_ – see Figure 1A of Gutenkunst et al. [56]. In the particular case of Equation (5.6), *σ*_2_ = 0 the major axis has infinite length and the ellipse degenerates to two parallel lines. This two-parameter case is illustrated in Figure 5 for both *Q* = 1 and *Q*_*∞*_ = 1.

**Figure 5.**
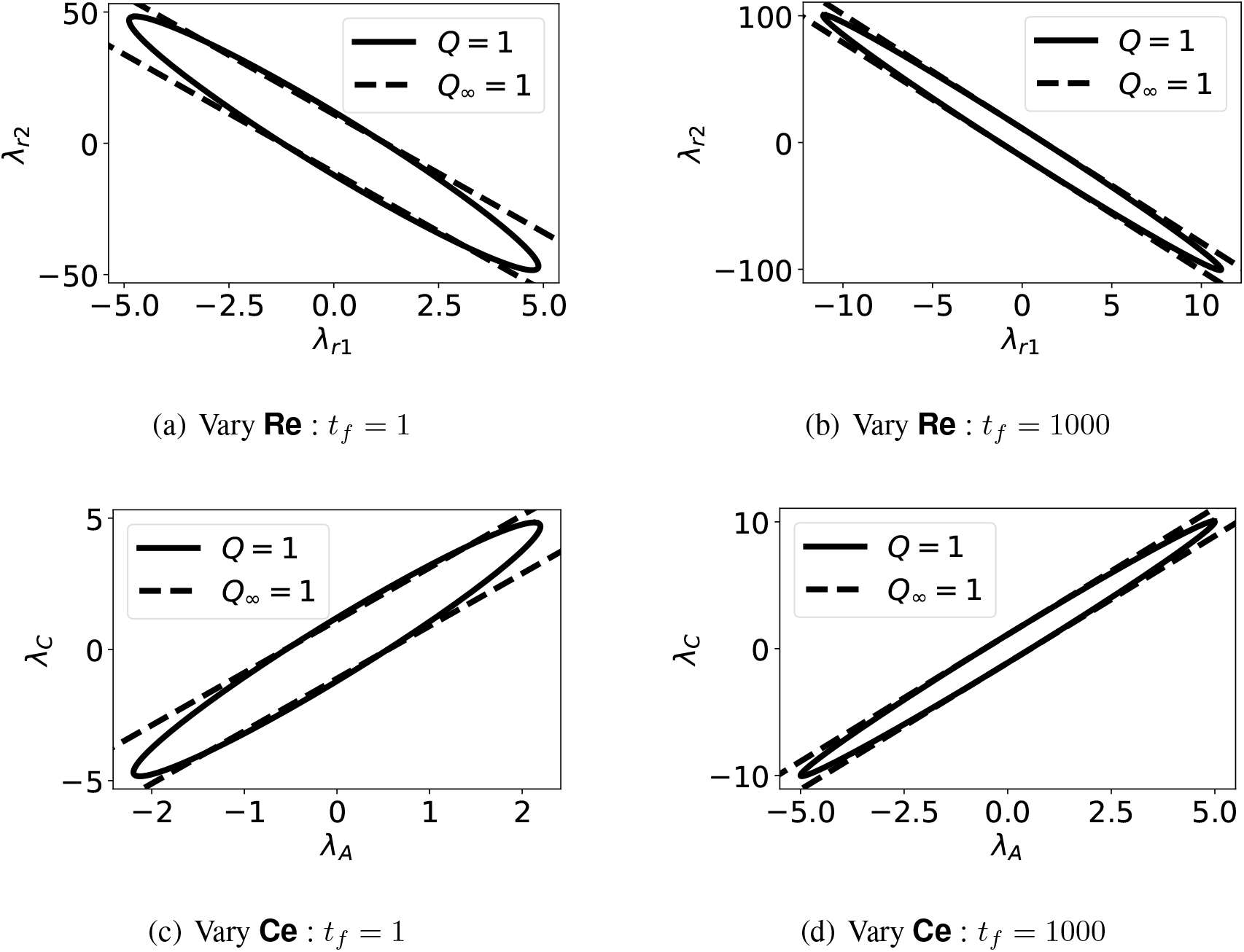
Sloppy parameters. As discussed in § 5.1, the contours of the two-parameter cost function corresponding to constant values *Q* = 1 are ellipses in parameter space whereas *Q*_*∞*_ = 1 corresponds to two parallel lines. (a) The parameters *λ*_*r*1_ and *λ*_*r*2_, corresponding to the two **Re** components, are varied. (b) As (a) but with a long timescale; as expected, *Q ≈ Q*_*∞*_. (c) The parameters *λ*_*A*_ and *λ*_*C*_, corresponding to the two chemostat **Ce** components, are varied. (d) As (c) but with a long timescale.

### 5.2 Example A ⇄B⇄ C

Here, we consider the steady-state sensitivity of the biochemical pathway A ⇄B ⇄C. For simplicity, consider the particular case where only the two sensitivity parameters λ_1_ and λ_2_ (corresponding to **Re**:**r**_**1**_ and **Re**:**r**_**2**_) are varied and the flow through **Re**:**r**_**1**_ is measured. The case where only the two sensitivity parameters *λ*_*A*_ and *λ*_*C*_ (corresponding to **Ce**:**A** and **Ce**:**B**) are varied can be analysed in a similar fashion.

The eigenvalue *σ*_1_ and eigenvector *V*_1_ corresponding to *H*_*∞*_ are given by:

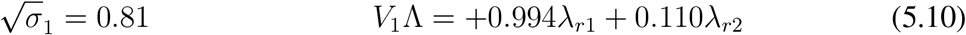

When *t*_*f*_ = 1, the two eigenvalues and eigenvectors of *H* are given by:

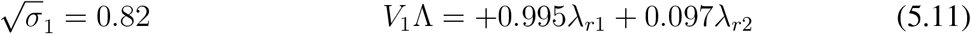

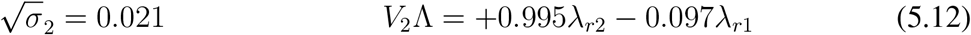

The ellipse corresponding to *Q* = 1 (5.2) appears in Figure 5(a) together with the two parallel lines corresponding to *Q*_*∞*_ = 1 (5.9).

When *t*_*f*_ = 1000, the two eigenvalues and eigenvectors of *H* are given by:

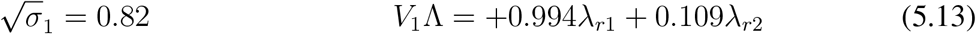

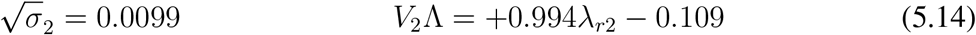

In this case, the first eigenvalue and eigenvector of *H* are close to that of *H*_*∞*_; and the second eigenvector *σ*_2_ *≈* 0. Thus, in this case, *H ≈ H*_*∞*_. The ellipse corresponding to *Q* (5.2) appears in Figure 5(b) together with the two parallel lines corresponding to *Q*_*∞*_ (5.9).

## 6 Example: Modulated Enzyme-catalysed reaction

Enzyme-catalysed reactions are an important control mechanism in biochemical systems [66]. One such mechanism is an enzyme-catalysed reaction with competitive activation and inhibition; which has been analysed within the bond graph context [53] and is shown in Figure 6. This simple system is used to illustrate the sensitivity methods of this paper. In particular, the dynamic sensitivities are derived and compared with the steady-state gain *g*_*∞*_ and time constant *τ* characterisation of the linear response. In addition, the variation of steady-state gain *g*_*∞*_ with steady-state flows through **Re**:**r**_**1**_ and **Re**:**r**_**2**_ is investigated.

**Figure 6.**
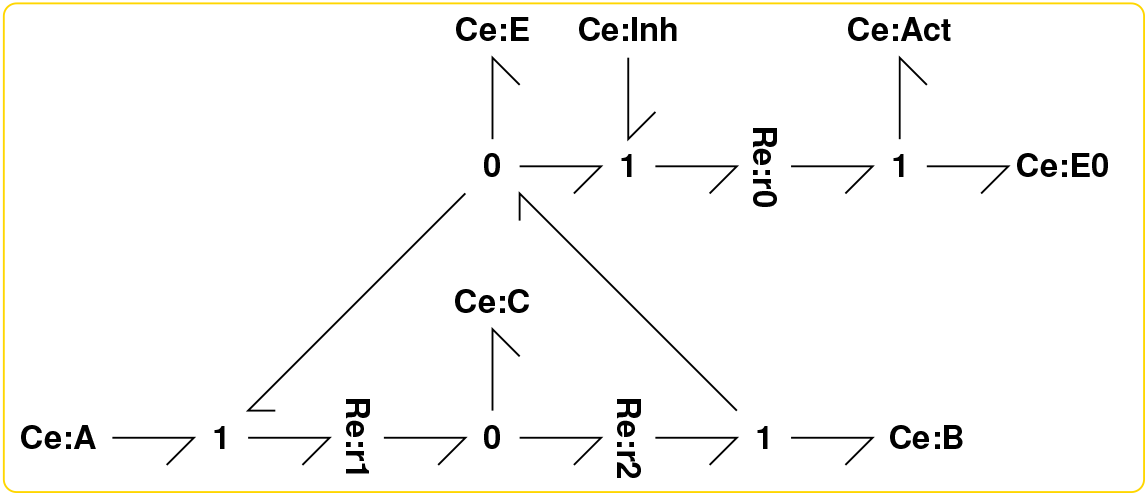
Modulated Enzyme-catalysed reaction: For the purposes of this example, all parameters are unity and all initial states are unity *except* the total enzyme amount e_0_ = ***x***_*E*_ +***x***_*C*_ +***x***_*E*0_ = 10 and ***x***_*B*_ = 10^−3^.

The four plots of Figure 7 show the dynamic sensitivity of the normalised flows in **Re**:**r**_**1**_ and **Re**:**r**_**2**_ to step changes in 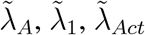 and 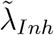. In the case of 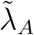 and 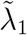, the flow through **Re**:**r**_**1**_ changes instantaneously and that through **Re**:**r**_**2**_ has a steady-state gain of about *g*_*∞*_ = 1.60 for 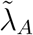 and *g*_*∞*_ = 0.80 for 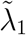. The normalised time constant is about *τ* = 0.32 in each case. In the case of 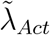 and 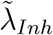, the flow through both **Re**:**r**_**1**_ and**Re**:**r**_**2**_ does not change instantly and has has a steady-state gain of about *g*_*∞*_ = *±* 0.8 and time constant of about *τ* = 0.4.

**Figure 7.**
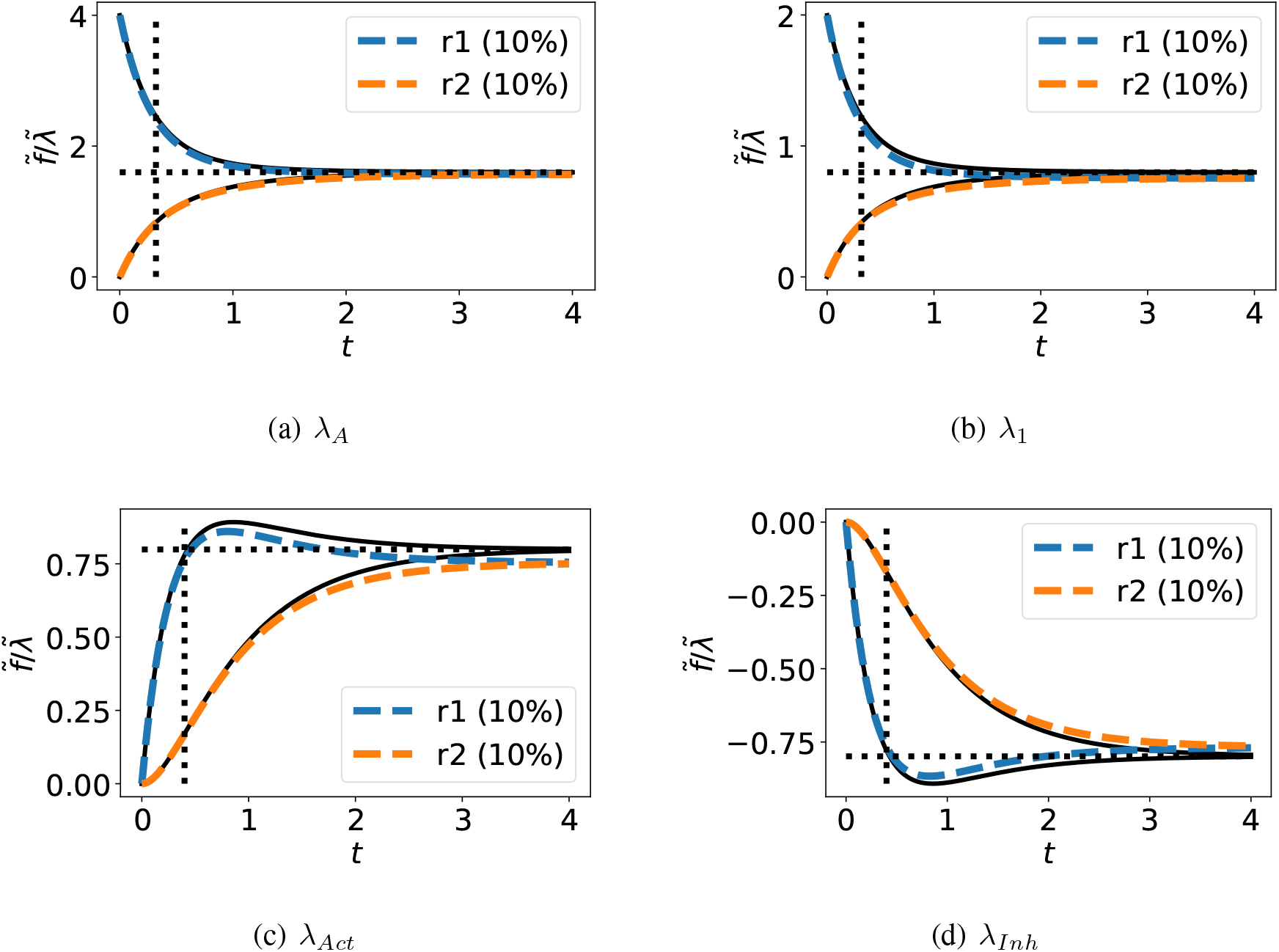
Modulated enzyme-catalysed reaction: Dynamical Sensitivities. The normalised temporal responses of the flows through **Re**:**r1**and **Re**:**r2** to perturbations in (a) ***K***_*A*_; (b) ***κ***_1_; (c) ***K***_*Act*_; (d) ***K***_*Inh*_ represented by the corresponding perturbation parameters *λ*_*A*_, *λ*_1_, *λ*_*Act*_ and *λ*_*Inh*_ respectively. Parameters are perturbed using a 10% step change in the perturbation parameter *λ*. The black line is the response of the linearised sensitivity system to the step change in the perturbation parameter *λ*, the horizontal dashed black line the corresponding DC gain *g*_*∞*_ and the vertical dashed black line the corresponding time-constant *τ*. The coloured dashed lines show the response of the simulated (nonlinear) sensitivity system to the same perturbation in *λ*.

Because the dynamics of the enzyme-catalysed reaction with competitive activation and inhibition are non-linear, the linearised gains are a function of the operating point. Figure 8(a) shows how the steady-state flow 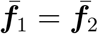 varies with the amount of substrate ***x***_*A*_; this is the typical reversible Michaelis-Menten curve saturating at a value dependent on the total enzyme. The system steady-state state is computed for a range of flows and the corresponding linearised gains are plotted in Figures 8(b) – 8(d). At small flows, the steady-state gains g_*∞*_ in Figures 8(b) – 8(d) are small because they correspond to flow sensitivity. At high flows, the flow is constrained by the maximum Michaelis-Menten flow *V*_*max*_ = *e*_0_κ_2_*K*_*c*_ where *e*_0_ is the total amount of bound and unbound enzyme; and so is independent of all of the sensitivity parameters shown except for *κ*_2_. Hence *g*_*∞*_ = 0 when the flow is saturated in all cases shown except for κ_2_.

**Figure 8.**
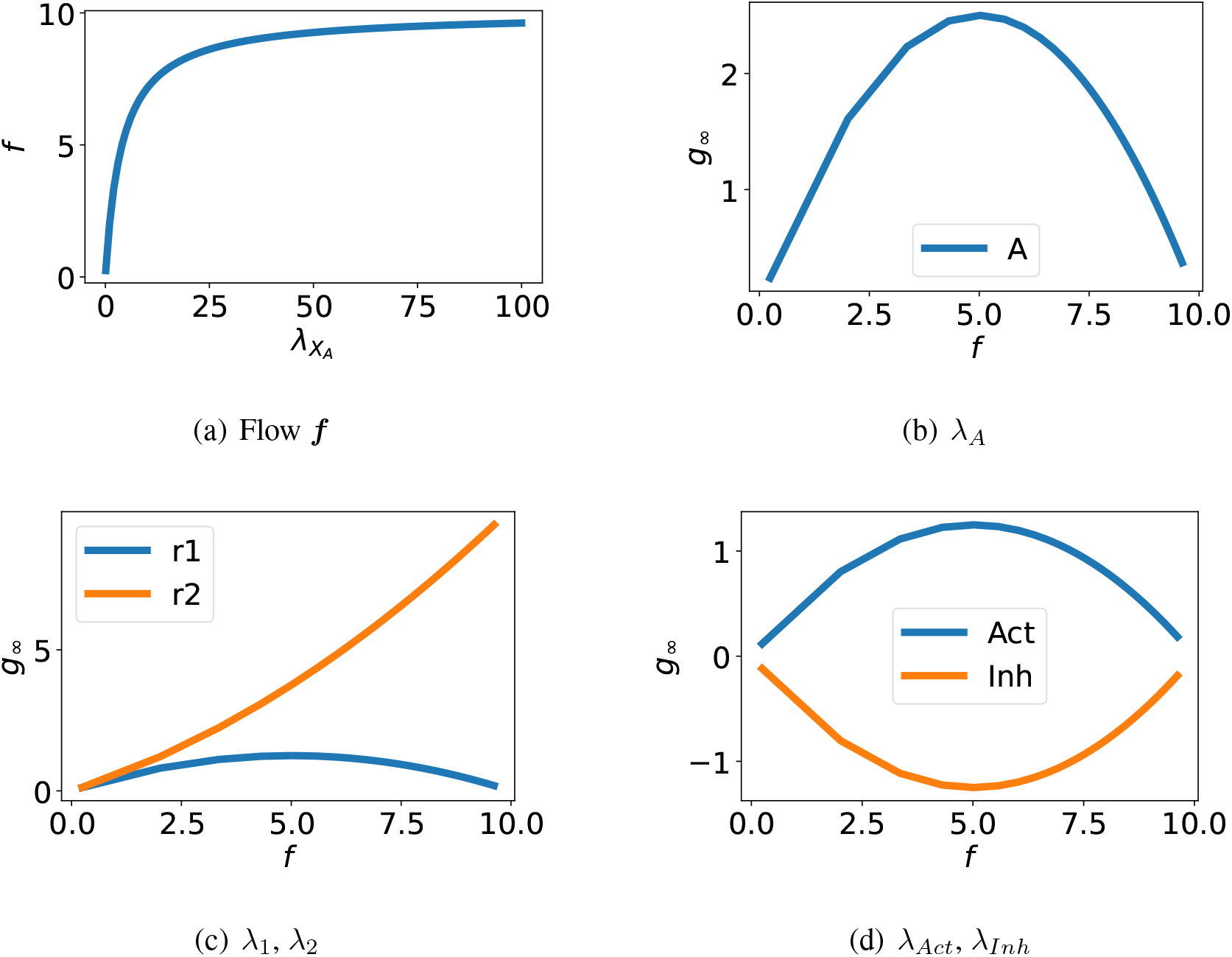
Modulated enzyme-catalysed reaction: Variation of sensitivity with steady-state. (a) The steady-state flow ***f*** = ***f***_1_ = ***f***_2_ is a non-linear function of the substrate concentration: the classical Michaelis-Menten curve. The corresponding steady-state values of the state of each **Ce** component is computed for a number of values of ***f***. (b) The sensitivity gain relating flow to variation in *λ*_*A*_ is plotted against the steady-state flow ***f***. (c) As (b) but with parameters *λ*_1_ and *λ*_2_ corresponding to **Re**:**r1** and **Re**:**r2**. (c) As (b) but with parameters *λ*_*Act*_ and *λ*_*Inh*_ corresponding to **Ce**:**Act** and **Ce**:**Inh**. As expected, activation increases, and inhibition decreases, steady-state flow; in both cases the modulation is most effective at flows corresponding to half the saturated flow.

## 7 Example: Pentose Phosphate Pathway

The Pentose Phosphate Pathway [67, 68] provides a number of different products from the metabolism of glucose. This flexibility is adopted by proliferating cells, such as those associated with cancer, to adapt to changing requirements of biomass and energy production [69, 70]. The

*E. coli* Core Model [71, 72] is a well-documented and readily-available stoichiometric model of a biomolecular system. As discussed previously [45], species, reactions and stoichiometric matrix pertaining to the Pentose Phosphate Pathway were extracted, a bond graph model created and parameters fitted using experimental data from *E. coli* [73]; the normalising time *t*_0_ = 7.95 s [45]. The reactions used in this model are listed in Appendix B.

This section examines the sensitivity properties of this model; in particular, § 7.1 looks at linearised sensitivity and why it is useful; § 7.2 looks at the linearisation error and § 7.3 examines the sensitivity from the sloppy-parameter viewpoint.

### 7.1 Linearised sensitivity

The sensitivity system was created using the approach of § 3.2 and linearised as discussed in § 2.4 and § 4. The advantage of linearisation is that steady-state gains *g*_*∞*_ and time-constants τ can be derived directly from the system bond graph model without the need for simulation or choice of perturbation magnitude. The errors involved in this simplification are discussed in § 7.2.

Figure 9 shows the steady-state gain *g*_*∞*_ from Equation (2.41) for the flows of three products: R_5_P, NADPH and G_3_P for two cases: sensitivity with respect to reaction constants and sensitivity with respect to chemostat states. Figure 10 shows the time constant *τ* for each of the six cases of Figure 9 computed as discussed in § 2.4.

**Figure 9.**
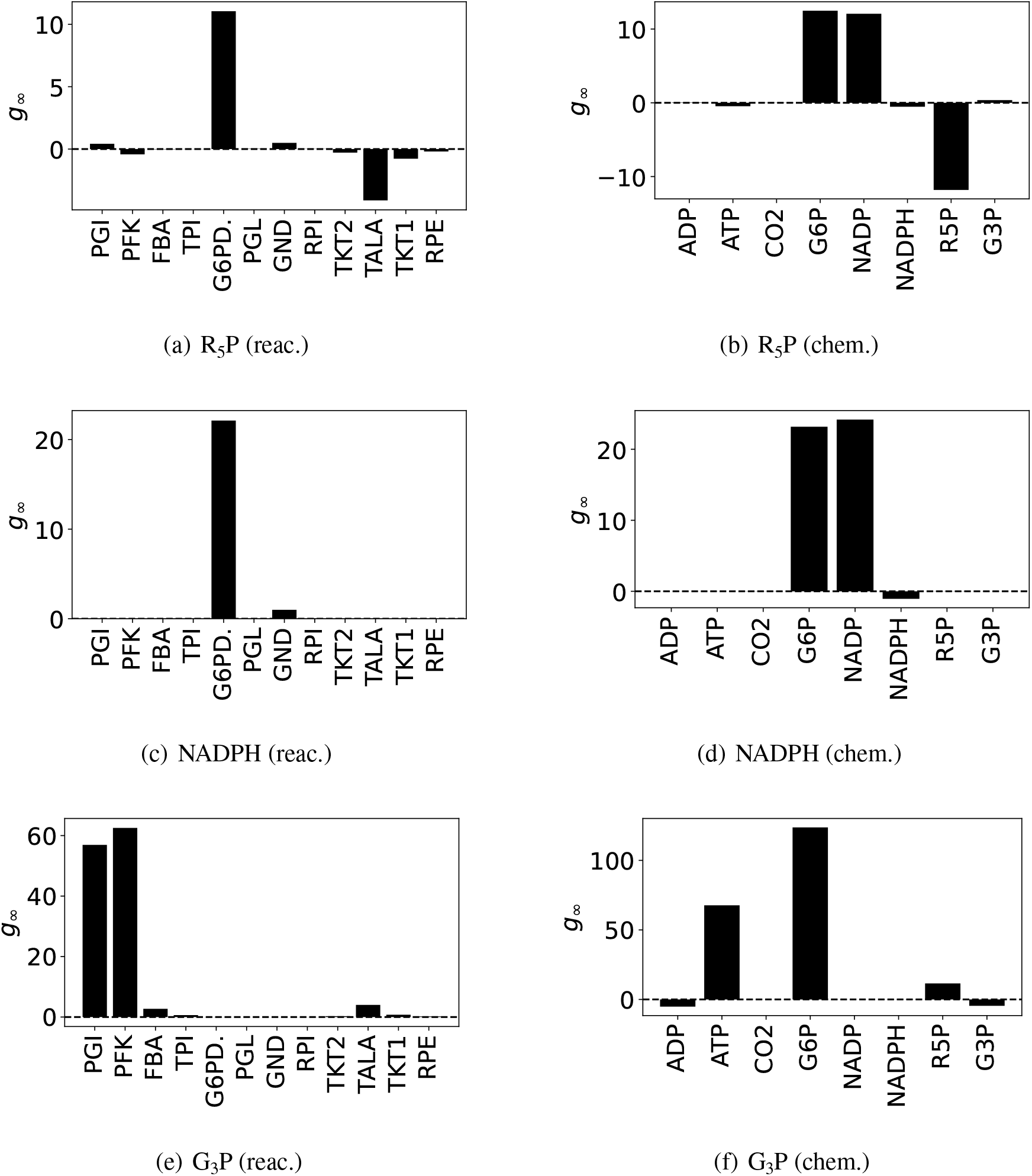
Pentose Phosphate Pathway: Steady-state sensitivity *g*_*∞*_. (a) The steady-state sensitivity *g*_*∞*_ of the flow of product R_5_P with respect to the reaction sensitivity variables *λ*_**Re**_ are shown for all reactions in the network. The dominant reaction is G_6_PDH_2_R. (b) The steady-state sensitivity *g*_*∞*_ of the flow of product R_5_P with respect to chemostat sensitivity variables *λ*_**Ce**_. Flow is increased primarily by G_6_P and NADP; flow is decreased primarily by R_5_P. (c) As (a) but for product NADPH; again, the dominant reaction is G_6_PDH_2_R. (d) As (b) but for product NADPH. Again, flow is increased primarily by G_6_P and NADP. (e) As (a) but for product G_3_P; in this case, the dominant reactions are PGI and PFK. (f) As (b) but for product G_3_P; in this case flow is increased primarily by G_6_P and ATP.

**Figure 10.**
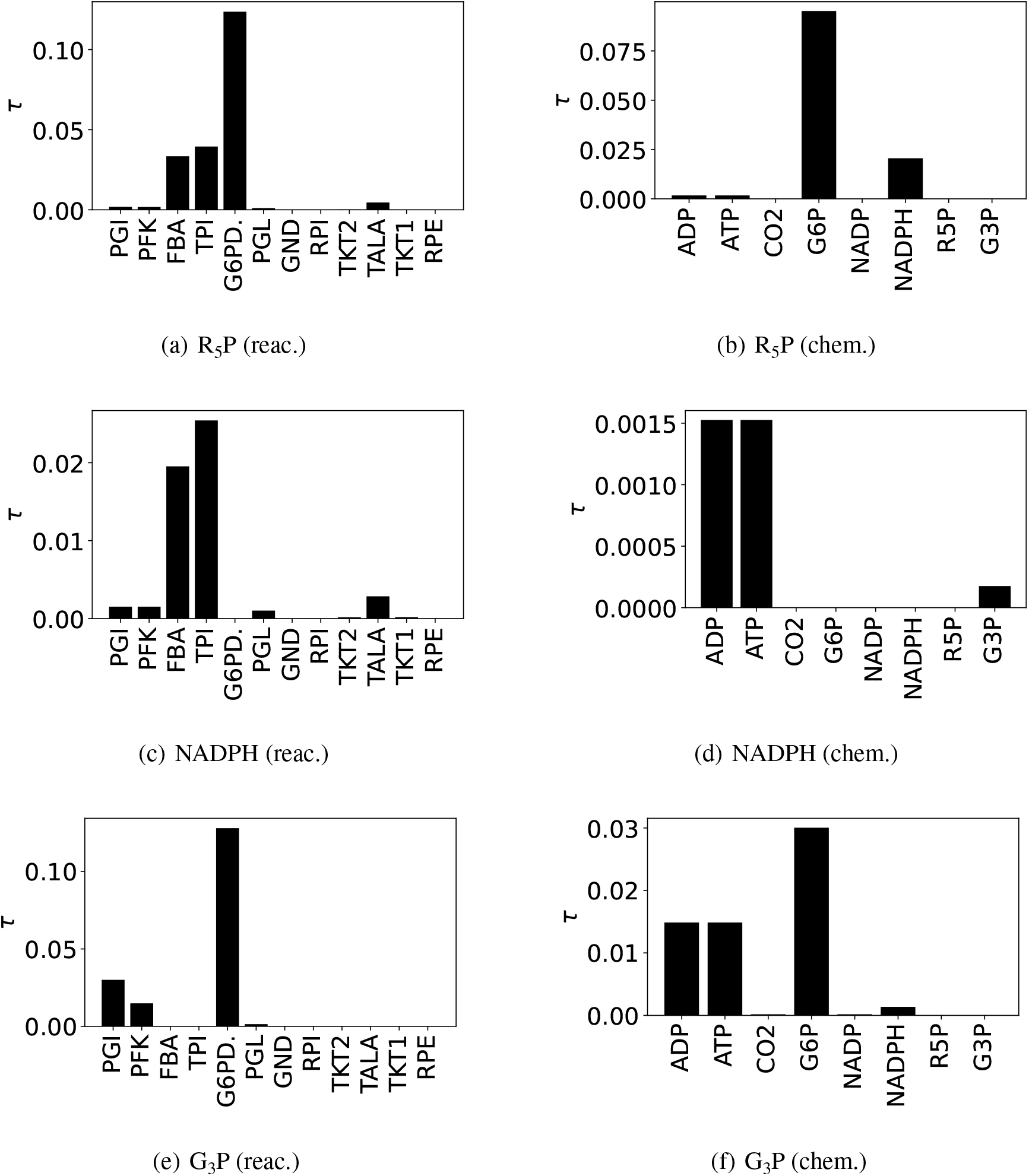
Pentose Phosphate Pathway: Sensitivity normalised time constant *τ*. The normalising time *t*_0_ = 7.95 s [45]. (a) The Sensitivity time constants *τ* of the flow of product R_5_P with respect to the reaction sensitivity variables *λ*_**Re**_ are shown for all reactions in the network. The sensitivity time constant for the dominant reaction G_6_PDH_2_R is about *τ* = 0.1 – see Figure 11. (b) The Sensitivity time constant *τ* of the flow of product R_5_P with respect to chemostat sensitivity variables *λ*_**Ce**_. The time constant for G_6_P is greater than that for NADPH reflecting the greater number of intervening reactions. (c) As (a) but for product NADPH; the sensitivity to dominant reaction G_6_PDH_2_R has no delay (*τ* = 0) as NADPH is a product of that reaction. (d) As (b) but for product NADPH. The sensitivity to dominant species G_6_P has no delay (*τ* = 0) as NADPH is a product of the same reaction. (e) As(a) but for product G_3_P; the dominant reactions PGI and PFK have a relatively small *τ* due to their proximity to this product. (f) As (b) but for product G_3_P; the dominant substrates G_6_P and ATP have relatively small *τ* decreasing with proximity to the product.

The rate parameter associated with G_6_PDH_2_R has the most influence over R_5_P and NADPH production (Figure 9(a,c)). These rates of production are also most sensitive to the concentrations of G_6_P and NADP, and the concentration of R_5_P has a substantial effect on its own rate of production (Figure 9(b,d)). Different reactions and species are involved in the regulation of G_3_P production, with PGI and PFK being the primary reactions (Figure 9(e)); and G_6_P and ATP being the primary metabolites (Figure 9(f)).

### 7.2 Linearisation error

The linearised sensitivity system (§ 4) is an approximation to the nonlinear sensitivity system (§ 3.1) for two reasons: it varies with the steady state of the non-linear system about which the linearisation is performed and the discrepancy between the linear and non linear system responses increases with the amplitude of the parameter variation. The former is discussed in § 6, Figure 8; the latter is examined further in this section in the context of the Pentose Phosphate Pathway example.

Figure 11 shows the effect of varying the amplitude of the sensitivity variable λ_*G*6*PDH*2*R*_. Figure 11(a) gives the non-linear deviation of the flow Δ***f***_*R*5*P*_ from its steady-state value normalised by the change in sensitivity variable Δλ_*G*6*PDH*2*R*_ for three values of the sensitivity vari-able Δ*λ*_*G*6*PDH*2*R*_. As expected, the error increases with Δ*λ*_*G*6*PDH*2*R*_ but, in this case, a tenfold increase only gives a 20% error. Figure 11(b) gives a similar result for product NADPH and Figure 11(c) gives a similar result for substrate G_6_P. Figure 11(d) summarises these results in terms of the steady-state linearisation error *ϵ* where:

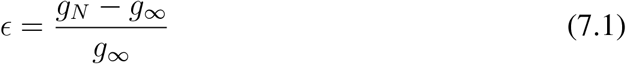

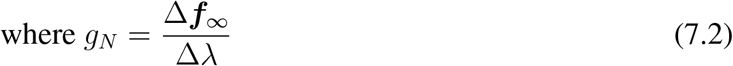

**Figure 11.**
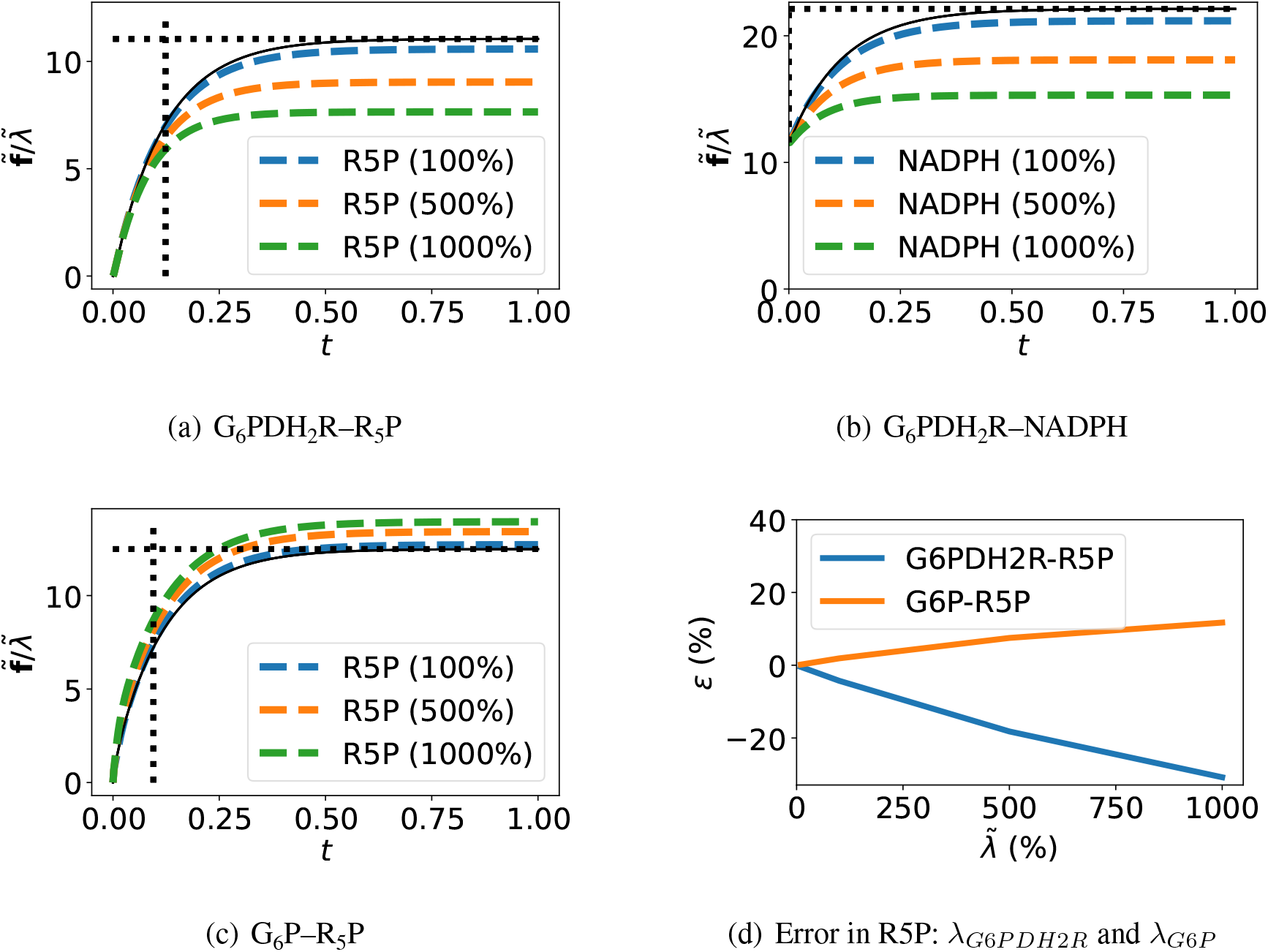
Sensitivity approximation. (a) The flow of product R_5_P is shown for a 100%, 500% and 1000% change in parameter *λ*_*G*6*PDH*2*R*_. (b) As (a) but showing flow of product NADPH (c) As (a) but with change in the concentration of G_6_P. (d) Linearisation error. *ϵ* = (*g*_*N*_ *− g*_*∞*_)/*g*_*∞*_ where *g*_*∞*_ is the steady-state (DC) gain of the linearised system and 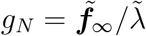 where 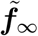 is the steady-state value of 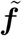 in (a) and (c). The gains are shown for two cases: sensitivity of R_5_P with respect to G_6_PDH_2_R and G_6_P.

### 7.3 Sloppy parameters

This section applies the methods of § 5 to the Pentose Phosphate Pathway model. In particular, the chemostat flows associated with product R_5_P is examined. Because this is a high-order system, the eigenvalue/eigenvector approach can be utilised to simplify the results by discarding small eigenvalues and small components of eigenvectors. In this case eigenvalues less than 1% of the largest eigenvalue are discarded, and within each eigenvector, components less than 10% of the largest element are discarded. For illustration, the sensitivity of the flow of R_5_P is investigated as all reaction constants are varied. In each case, the results using the *Q*_*∞*_ cost function in Equation (5.9) are compared with those from *Q* (5.2) when the final time ***t***_*f*_ = 0.82 is chosen by the `step_response` function of the Python Control Systems Library [74]).

The eigenvalue σ_1_ and eigenvector *V*_1_ corresponding to *H*_*∞*_ are given by:

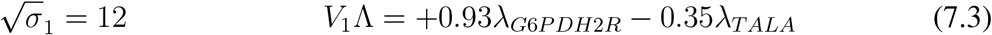

When multiplied by 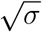, the eigenvector elements correspond to the two largest values of *g*_*∞*_ given in Figure 9(a). This illustrates the point discussed in § 5: sloppy analysis of the linearised response of the sensitivity system using *H*_*∞*_ gives the same information as *g*_*∞*_ obtained without simulation.

The significant eigenvalues, and corresponding significant elements of the eigenvector corresponding to *H* are given by:

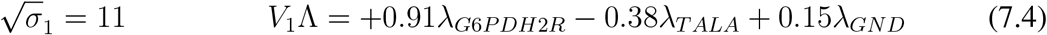

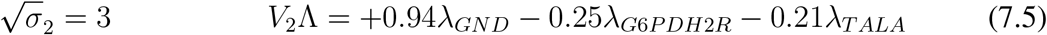

As discussed in § 5, *H*_*∞*_ corresponds to an infinite time span whereas *H* corresponds to the length of the simulation, in this case ***t***_*f*_ = 0.82. Hence *H* is dependent on the initial, transient, part of the response whereas *H*_*∞*_ only depends on the steady-state response. The fact that GND appears in the eigenvectors of *H*, but not in the eigenvectors of *H*_*∞*_, implies that perturbation of GND affects the initial response but has no significant effect on the steady-state response. Indeed, the corresponding unit step response initially rises to over 10 before falling back to the steady-state value of *g*_*∞*_ *≈* 0.5. This behaviour is consistent with experimental evidence; in particular, whilst GND is not seen as the major rate-controlling step of the Pentose Phosphate Pathway [75], studies have proposed its involvement in the short-term response to oxidative stress [76].

## 8 Conclusion

It has been shown that the sensitivity properties of the model of a biochemical system modelled using the bond graph formulation can be examined by creating a corresponding sensitivity bond graph model. The sensitivity model can be created by either replacing bond graph components with variable parameters by a corresponding sensitivity component or via a stoichiometric approach. Either approach can be used to automatically convert the original bond graph to a sensitivity bond graph. Within the sensitivity model, the parameter variation appears as a set of additional inputs modelled as bond graph chemostat components. Hence previously developed linearisation approaches can be used to linearise the nonlinear sensitivity system with respect to parameter variation to give local sensitivity results.

The well known “sloppy parameter” method, and its corresponding cost-functions have been incorporated into the sensitivity approach of this paper and lead to illuminated eigenvalue/eigen-vector decomposition of a Hessian matrix associated with the local sensitivity results.

The results have been illustrated using a previously-derived model of the Pentose Phosphate Pathway. In this particular case, the linearised sensitivity system is compared to the nonlinear case and found to form an accurate approximation. As is well-known [67, 22.6], the reaction G_6_PDH_2_R is an important regulator of the Pentose Phosphate Pathway; the sensitivity analysis does indeed show that product flows depend strongly on this reaction.

The methods presented here could potentially be used to analyse omics data. While statistical and bioinformatics methods can reveal analytes of interest, conducting a sensitivity analysis on a mechanistic model can help researchers to assess the importance of potential changes in these analytes. Recent metabolomics and proteomics analysis of heart tissue [77] gives experimental evidence for how disease affects the heart in terms of its biochemistry. Combining this differential omics analysis with corresponding biochemical sensitivity models will allow us to map the omics data onto medically-significant physical phenomena such as cardiovascular disease. Parameters with high sensitivity may point towards potential targets for biomedical interventions.

The bond graph sensitivity system replaces parameter variation by inputs. In this paper, parameter sensitivity is examined by setting these inputs to a fixed value; future work will examine time-varying parameters as modelled by time-varying inputs to the sensitivity system.

A major challenge in systems biology is model parameterisation, particularly for large-scale models [78]. Sensitivity analysis can inform experimental design to reduce uncertainty in model predictions. Our results in § 7.2 indicate that not all parameters are equally important for the behaviour of a model, and that different parameters are important for describing different aspects of model behaviour. This observation has been mirrored in other sensitivity studies [56, 79]. Thus, determining the ‘stiff’ combinations of parameters can help to optimise for knowledge gained with limited experimental resources. Furthermore, analysing the time constants can be used to assess the biological relevance of parameter sets in machine learning approaches to parameterisation [80, 81]. Our approach builds on existing techniques by reframing parameters in a thermodynamically consistent context, which can in some cases reduce the number of parameters and better distinguish between the individual contributions of reactions and metabolites [82, 83]. There is a wealth of control-theoretic results applicable to linear systems. As indicated in the example of § 6, modulating the parameters of enzyme-catalysed reactions is a key strategy in human cellular control systems. The sensitivity models of this paper provide a foundation for applying such control-theoretic results to understand biochemical control systems. These methods are particularly relevant for attempts to rationally engineer pathways in synthetic biology, where sensitive parameters indicate potential targets of modification [84]. In many cases, biological networks are designed to be robust to noise, independent of parameter values [85, 86]. However, this robustness often comes at a cost to resource and energy consumption [41, 87], and in some cases, noise is harnessed for biological control [88, 89]. A bond graph modelling approach frames these behaviours in a thermodynamic context, allowing investigations into trade-offs between energy consumption and performance.

## Author Contributions

PJG prepared the numerical results and drafted the manuscript; PJG and MP discussed the results and revised the manuscript. PJG and MP approved the final version for publication. The authors would like to thank the referees for their constructive comments which have enhanced the paper.

## Acknowledgements

PJG would like to thank the Faculty of Engineering and Information Technology, University of Melbourne, for its support via a Professorial Fellowship. MP was supported by a Postdoctoral Research Fellowship from the School of Mathematics and Statistics at the University of Melbourne.

## Data Accessibility

The figures and tables in this paper were generated using the Jupyter notebooks and Python code available at https://github.com/gawthrop/Sensitivity23.

## A Example A ⇆ B ⇆ C: numerical calculations

In this special case, equation (3.13) becomes:

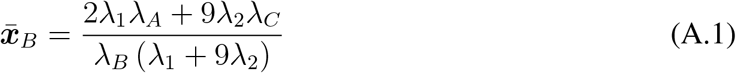

When all parameters are at their nominal values so that each *λ* = 1:

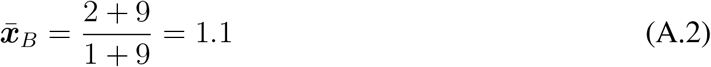

Similarly in the special case, the steady state values of ***f***_1_ and ***f***_2_ are

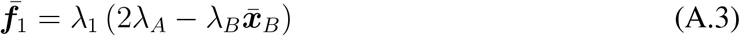

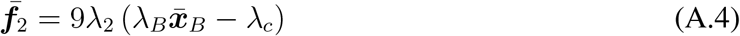

and when all parameters are at their nominal values:

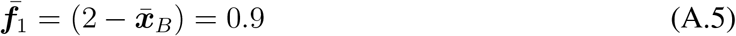

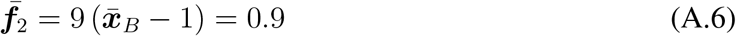

The steady-state sensitivities can be computed from Equations (A.1) – (A.4) by taking derivatives and substituting unit values for the remaining λ; for example:

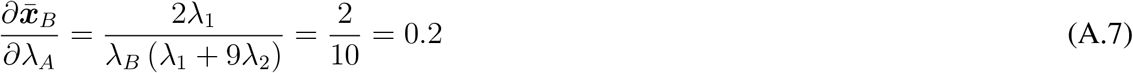

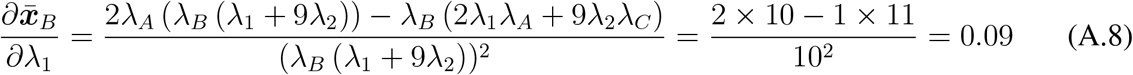

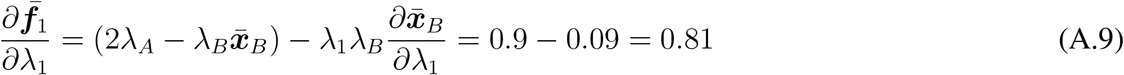

## B Reactions of the Pentose Phosphate Pathway

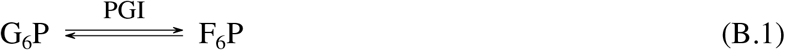

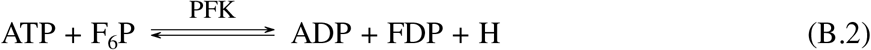

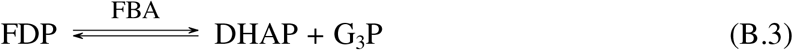

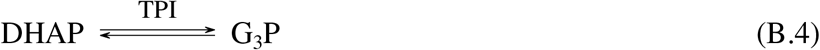

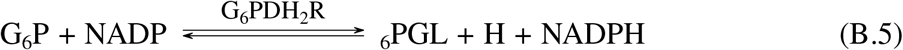

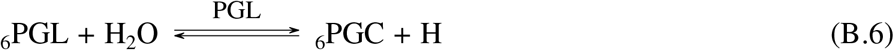

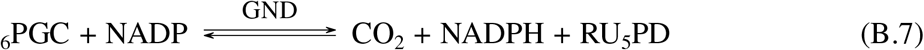

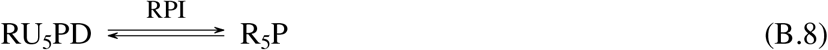

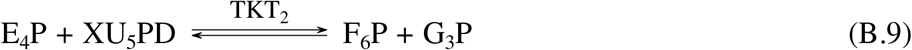

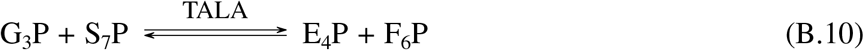

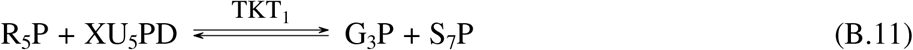

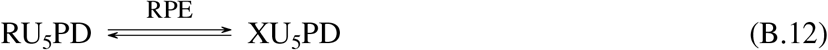

## Footnotes

1 Bold font is used to signify normalised variables.

